# Epithelial multicellular clustering enabled by polarized macrophages on soft matrices

**DOI:** 10.1101/2023.02.20.529258

**Authors:** Hannah Zmuda, Amit Pathak

## Abstract

Formation of epithelial structures of variegated geometries and sizes is essential for organogenesis, tumor growth, and wound repair. Although epithelial cells are predisposed with potential for multicellular clustering, it remains unclear whether immune cells and mechanical cues from their microenvironment influence this process. To explore this possibility, we co-cultured human mammary epithelial cells with pre-polarized macrophages on soft or stiff hydrogels. In the presence of M1 (proinflammatory) macrophages on soft matrices, epithelial cells migrated faster and subsequently formed larger multicellular clusters, compared to co-cultures with M0 (unpolarized) or M2 (anti-inflammatory) macrophages. By contrast, stiff extracellular matrix (ECM) disabled active clustering of epithelial cells due to their enhanced migration and cell-ECM adhesion, regardless of macrophage polarization. We found that the co-presence of soft matrices and M1 macrophages reduced focal adhesions, but enhanced fibronectin deposition and non-muscle myosin-IIA expression, which altogether optimize conditions for epithelial clustering. Upon Rho-associated kinase (ROCK) inhibition, epithelial clustering was abrogated, indicating a requirement for optimized cellular forces. In these co-cultures, Tumor Necrosis Factor (TNF)-α secretion was the highest with M1 macrophages and Transforming growth factor (TGF)-β secretion was exclusively detectable in case of M2 macrophages on soft gels, which indicated potential role of macrophage secreted factors in the observed epithelial clustering. Indeed, exogenous addition of TGB-β promoted epithelial clustering with M1 co-culture on soft gels. According to our findings, optimization of both mechanical and immune factors can tune epithelial clustering responses, which could have implications in tumor growth, fibrosis, and would healing.

**Summary:** Authors show proinflammatory macrophages on soft matrices enable epithelial cells to form multicellular clusters. This phenomenon is disabled on stiff matrices due to increased stability of focal adhesions. Inflammatory cytokine secretion is macrophage-dependent, and external addition of cytokines accentuates epithelial clustering on soft matrices.

**Impact Statement:** Formation of multicellular epithelial structures is critical to tissue homeostasis. However, it has not been shown how the immune system and mechanical environment affect these structures. The present work illustrates how macrophage type affects epithelial clustering in soft and stiff matrix environments.

## Introduction

Epithelial cell clustering and formation of complex multicellular structures are critical for fundamental biological process including development, wound healing, and malignant diseases (1). For epithelial cells to maintain and enable such physical functions of morphogenesis, a balance of intra- and intercellular forces is required (2). In collective epithelial migration, cells coordinate with their neighbors to spread on mechanically distinct underlying substrates and yet stick together via adherens junctions (3). One important physical function of adherens junctions is to transmit actomyosin contractile forces generated within single cells across cell-cell junctions and thus regulate macroscale tissue folding and complex shape formation (3, 4). Cells generate forces through mechanotransductive signaling due to Rho GTPases (5, 6) and their feedback with focal adhesions and actomyosin motors driven by crossbridge cycling of non-muscle myosin IIA (NM2A) on filamentous actin (F-actin) fibers (7). Cell level forces coordinate with other cells and ECMs to enable epithelial cluster formation and tissue compaction, which is of functional importance in wound healing and tumor growth that also feature a complex immune system consisting of macrophages of varying polarities (8–10). Although recent work has shown that elasticity and viscoelasticity of extracellular matrices (ECMs) affect epithelial migration and clustering (2, 11), it remains unexplored whether the co-presence of macrophages of defined polarization also influences such epithelial responses.

During physiological processes, as in development, cells undergo mechanical and chemical changes to undergo tissue shape change. One such change is the aggregation of cells, such as in mammalian embryogenesis, organogenesis, or carcinomas. Tissue compaction mechanics can be related to the physics of wetting, such that competition of cell-cell adhesions and surface tension (cell tractions and contractility) occurs between neighboring cells. According to previous work, breast cancer cells on hydrogels collectively contract and form multicellular aggregates due to increased tractions, intracellular tension, and simultaneous rise in E-cadherin (E-cad) expression (12). During tissue retraction, also known as ‘dewetting’, both E-cad and phosphorylated myosin light chain (pMLC) are highly expressed, the former recruiting the latter (12, 13). Moreover, a combination of weakened cell-ECM adhesions, and NM2A-dependent contractile forces allow cells to form cell aggregates on soft, but not on stiff, hydrogels (11). In addition to matrix stiffness, changes in the type of extracellular protein ligands, e.g., collagen I, collagen IV, laminin, or fibronectin, also influence epithelial responses (14, 15). Although there is emerging understanding of how cells coordinate their forces and adhesions with one another and with the ECM, it is less explored how epithelial aggregation is influenced by the presence of other dissimilar cells.

Multiple cell types are involved for tissue bodies to properly form and repair. Additional cells can include fibroblasts, endothelial cells, and immune cells. Among these, macrophages, the scavenger immune cells, are known to clear bacteria and foreign body substances through phagocytosis and can both assist and kill tumor cells in cancer progress according to macrophage polarization (9, 10). During development of *Drosophila*, macrophages must disseminate into the germ band by squeezing between ectodermal cells during cell division, facilitated by focal adhesion disassembly (16). Additional work investigating the co-presence of macrophages found endothelial cells spontaneously undergo capillary morphogenesis in the presence of cancer-associated macrophages due to upregulation of the adhesion molecule ICAM-1 (17). In response to chemical or biological activation, macrophages polarization can become either pro-inflammatory (M1) or anti-inflammatory (M2) (18, 19). These different polarizations enable macrophages to perform a variety functions – aid wound healing in M1 phenotype and assisting tumor progression by M2 macrophages (8, 18, 20, 21). Overall, previous work has shown that macrophages can cooperate or compete with different cell types to regulate tissue- and multicellular-scale processes.

In this study, we explore whether epithelial clustering can be optimized by the co-presence of macrophages of predefined polarization and stiffness of the underlying substrate. To regulate macrophage polarization, we directly activated THP-1, immortalized monocyte cell line, into M0, M1, or M2-like macrophages. We co-cultured these differentially polarized macrophages along with human mammary epithelial cells (MCF10A) atop soft (0.5kPa) or stiff (120kPa) polyacrylamide (PA) hydrogel substrates and studied epithelial responses. Indeed, we found that propensity for epithelial clustering increased with soft environment and pro-inflammatory (M1) macrophages. We also found that pharmacological perturbation of cellular mechanotransductive signaling disturbs the optimal balance of cellular forces and adhesion required for epithelial clustering. Overall, our results highlight a previously unappreciated role of the presence of macrophages along with soft environments in epithelial cluster formation, which is orchestrated by a balance of cellular forces and adhesions.

## Results

### Macrophage polarity regulates epithelial cluster formation on soft matrices

To study how epithelial cells respond to changes in macrophage polarization and matrix stiffness, we first cultured human monocytes directly on soft (0.5kPa) polyacrylamide (PA) gels coated with collagen-type I (Col1), which recapitulate soft tissue microenvironments better than the extremely stiff tissue culture plastic surface. We polarized these monocytes into unstimulated (M0), proinflammatory (M1), and anti-inflammatory (M2) macrophages (Figure 1A). We then seeded human mammary epithelial cells (MCF10A) with green fluorescing nuclei (GFP) atop the activated macrophages (Figure 1B). We observed MCF10A cells forming clusters within 40hr on soft gels (Movie 1 and 2; Figure 2A). We quantified temporal epithelial cell clustering by measuring the normalized intercellular distance (ID), calculated as distance between all pairs of nuclei within a frame overtime (Figure S1). As epithelial cells co-cultured with M0 or M1 macrophages begin to cluster, the ID value drops below one, indicating aggregation of nuclei relative to their initial position (Figure 2B). Cell viability is verified by the continued nuclear expression of GFP (Movie 1-3; Figure S2). As this epithelial clustering proceeded over time, we observed distinct differences in resulting shape and size of epithelial clusters after 60hr of live imaging (Figure S2, white arrow heads). Specifically, when co-cultured with M0 macrophages, epithelial clusters were many, sporadic, smaller in area, and rounder (Figure 2A.i). By contrast, in the presence of M1 macrophages, epithelial cells formed much larger and elongated clusters (Figure 2A.ii). However, epithelial cells with M2 macrophages rarely formed clusters (Movie 3). Overall, across multiple samples, we found that M1 macrophages enabled the formation of much larger epithelial clusters compared to M0 or M2 (Figure 2C).

**Figure 1.**
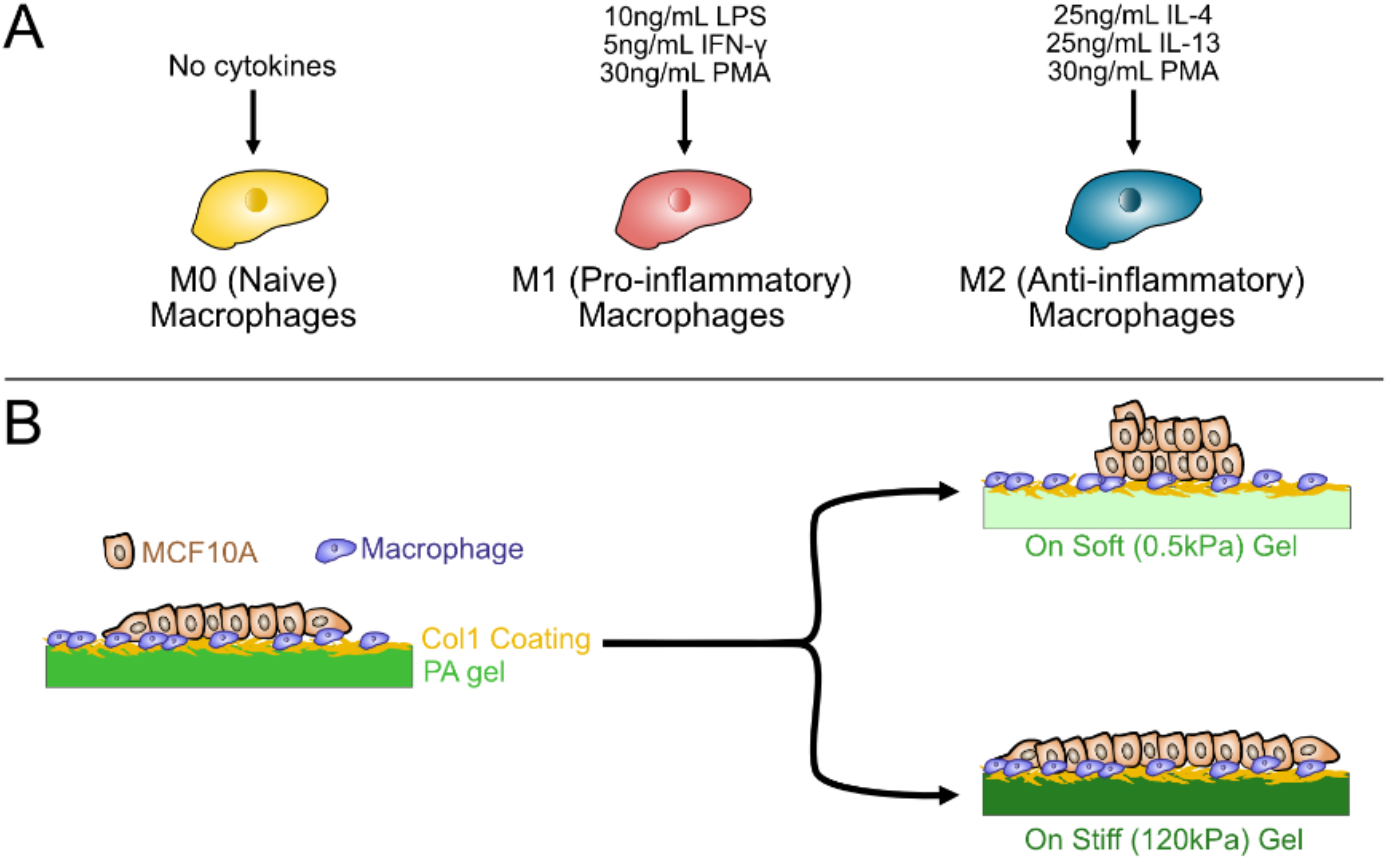
Experimental setup of epithelial cell co-culture with macrophages on hydrogels. **(A)** Immortalized monocytes were primed to macrophages by adding 30ng/mL PMA. The macrophages were then polarized in three ways: (1) M0 (naïve) without PMA, (2) M1 (proinflammatory) with 10ng/mL LPS, 5ng/mL IFN-*γ*, and 30ng/mL PMA, and (3) M2 (anti-inflammatory) with 25ng/mL IL-4, 25ng/mL IL-13, 30ng/mL PMA. **(B)** Macrophages directly activated on collagen type 1 (Col1)-coated soft or stiff polyacrylamide (PA) gels. After macrophage activation, human mammary epithelial cells (MCF10A) were seeded into a monolayer atop the macrophages.

**Figure 2.**
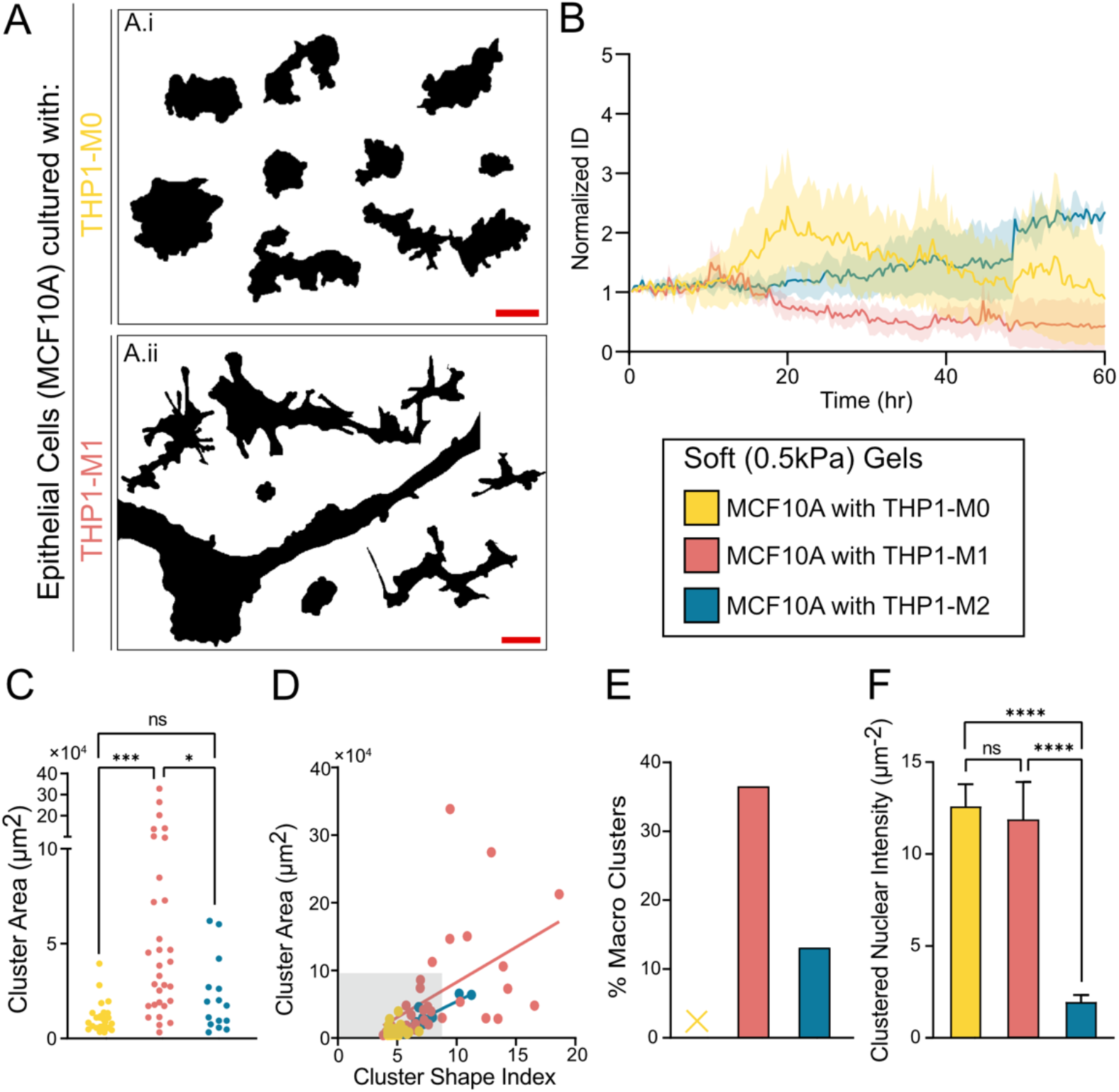
Epithelial cells cluster in different sizes according to macrophage type on soft matrix. **(A)** Outlines of epithelial clustering on soft PA gels when co-cultured with (A.i) M0-like macrophages and (A.ii) M1-like macrophages. Scale bar represents 100µm. **(B)** Line plot of the normalized intercellular distance (ID) represents the degree of clustering overtime. Epithelial cells in the co-presence of macrophages begin to cluster first, with their average going below 1 by t = 20hr. By the end of the experiment, both M0 and M1-like macrophages form cluster (Normalized ID < 1). **(C)** Area of individual clusters from different macrophage co-cultures. Larger epithelial cluster in M1 co-culture, compared to clusters with M0 and M2 macrophages. **(D)** Scatter plots comparing area and shape index of epithelial clusters with different macrophages. Lines represent linear regression of each macrophage type. **(E)** Percentage of epithelial clusters that fall outside of the gray box (area>10^5^µm^2^ and shape index>8), known as macro-clusters. “X” denotes no macro-clusters were formed outside of the gray box for the condition. **(F)** Bar graph representing the intensity of nuclei within clusters. Epithelial cells in M0 and M1 co-cultures clustered, creating clusters of nuclei and high fluorescence intensity. Error bars represent SEM. Data was analyzed using One-way ANOVA (with Tukey’s multiple comparison), with n = 3 biological replicates for all, to compare macrophage co-culture on soft PA gels.

To understand geometric differences in epithelial clusters, we plotted cluster area versus shape index (Figure 2D). Upon counting, we found over 35% clusters with M1 macrophages were in this category while 0% with M0 and <15% with M2 macrophages (Figure 2E). Moreover, nuclear intensity for clusters was higher in epithelial cells with co-presence of M0 and M1-like macrophages (Figure 2F). Based on the difference in cluster type, we defined epithelial ‘*macro-clusters*’ as those with both large area (>10^3^μm^2^) and shape index (>8) (plotted in Figure 2C). We categorized ‘*micro-clusters’* as the ones with smaller area and shape index, peppered randomly over the substrate. Although recent studies have shown multicellular clustering on soft substrates (12), our findings reveal that micro- and macro-clustering of distinct sizes can be enabled by the co-presence of macrophages of defined polarization.

### Macro-cluster formation of epithelial cells is associated with optimal migration phenotypes

Although it is well understood that cells can spontaneously form cell aggregates on soft matrices or in the presence of different ECM types (11, 14, 22, 23), it is not clear whether certain migration phenotypes are associated with potential multicellular clustering. Due to epithelial-mesenchymal transition (EMT), cell migration increases upon breakdown of microtissues and organoids (24). However, the converse is unclear – whether prior migration speed and order of individual cells within epithelial colony determine size and shape of ensuing clusters. To measure individual cell migration phenotypes leading up to collective clustering, we tracked nuclei of MCF10A cells overtime. When clustering occurs, cells collapse into the center of the cell colony (Movie 1-2). With M0 and M1 macrophages, we observed that epithelial cell clustering begins after approximately 20hr (Figure 3A.i and 3A.iii) and continues until clusters aggregate by ∼40hr (Figure 3A.ii and 3A.iv), which we refer to as ‘peak clustering time’ (20≤*t*≤40hr). We plotted velocity vectors directed towards net displacement and color-coded for average speed between the 20-40hr clustering time period noted above (Figure 3A). Using this spatial and temporal speed distribution, we observed higher speeds on cluster edges in M0 and M1 co-cultures (Figure 3A). In time-lapse videos, we observed much higher epithelial cell displacement with M1s (Movie 2) compared to other conditions. To better understand this qualitative observation, we plotted individual cell tracks between 20-40hr clustering time and found longer (>250µm) and more persistent epithelial cell trajectories with M1 macrophages, while cell tracks were shorter and more random with M0 or M2 macrophages (Figure 3B.i and 3B.iii, respectively).

**Figure 3.**
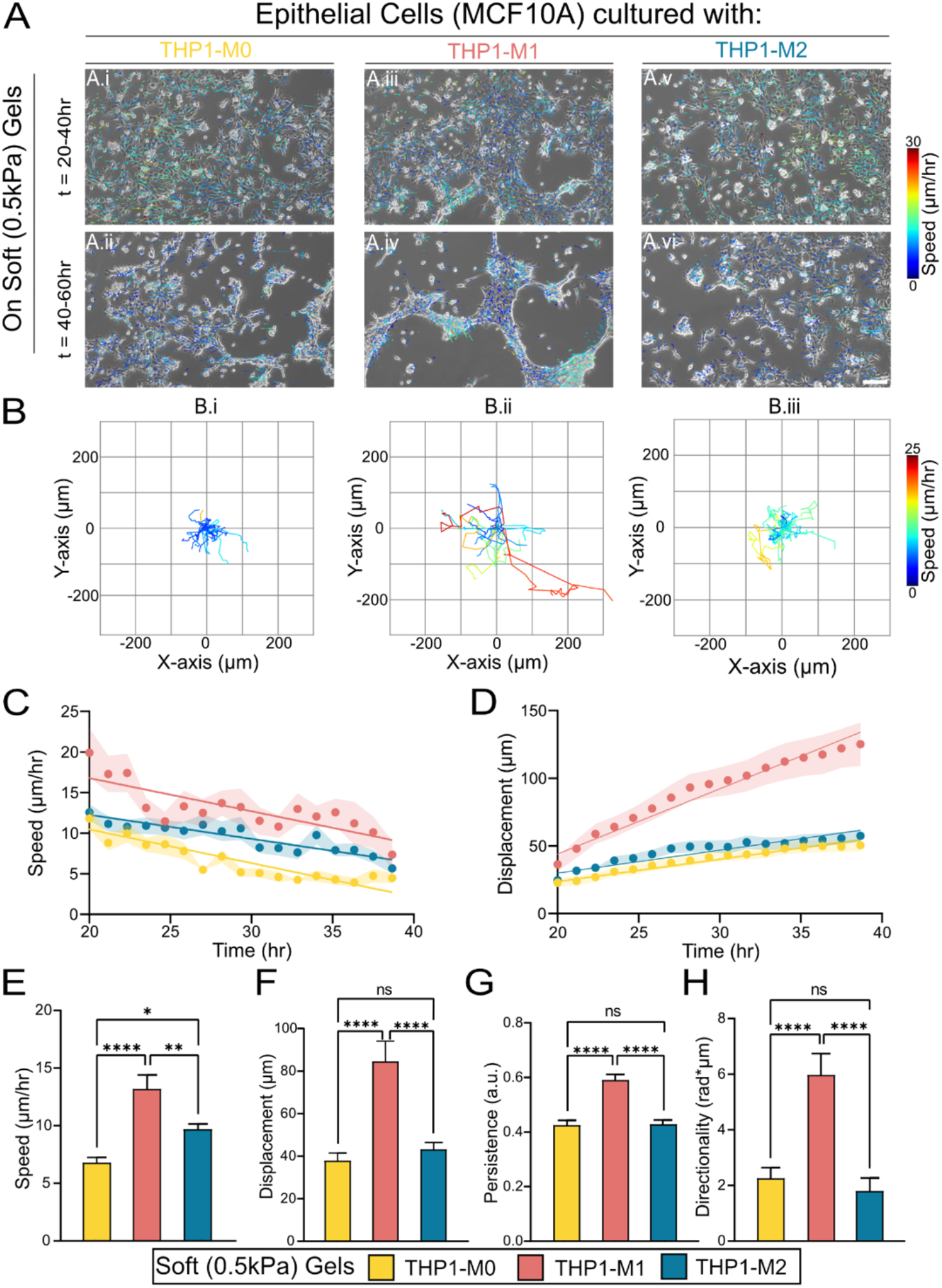
Higher cell migration precedes macro-clustering on soft matrices. **(A)** Representative quiver plots of epithelial tracks with overlayed brightfield images of MCF10A cells with (A.i, A.ii) M0-like, (A.iii, A.iv) M1-like, and (A.v, A.vi) M2-like macrophages at time *t*=20-40hr and *t*=40-60hr of the experiment, respectively. Quiver magnitudes represent cumulative distance of track during the time window. Quiver color represents average migration speed. **(B)** Representative plots of epithelial migration tracks (n = 25 cells). Tracks represent cumulative epithelial cell distance travelled between 20-40hr and color represents average migration speed for (B.i) M0, (B.ii) M1, and (B.iii) M2 co-cultures. (**C** and **D**) Cell migration speed and distance over time, based on manual tracking of cells migrating over 20-hour time window. Line represents linear regression; points are average. Shaded region represents standard error of the mean (SEM) from n≥3. **(E-H)** Bar plots represent average migration parameter with SEM from n≥3 replicates. Data was analyzed using One-way ANOVA (with Tukey’s multiple comparison) to compare macrophage co-culture on soft PA gels. ****P<0.0001, **P<0.01, *P<0.05, ns = not significant.

To further understand how cell migration temporally evolves during the clustering period, we plotted average instantaneous speed and net displacement of epithelial cells over time (Figure 3C-D). Consistent with video observations and individual cell tracks, speed and displacement of epithelial cells cultured with M1 macrophages were higher than those with M0 or M2 (Figure 3C). For all three macrophage types, epithelial cell speed reduced over time due to clustering, albeit of different sizes. However, the net displacement increased overtime because cells were moving farther from their initial position during clustering events (Figure 3D). Upon averaging over time and across samples, we found that migration speed, net displacement, linear persistence, and directionality of epithelial cells on soft matrices were significantly higher with M1 macrophages compared to their co-culture with either M0 or M2 macrophages (Figure 3E-H). Altogether, these migration trends indicate that faster and directionally persistent migration of epithelial cells is associated with their higher propensity for forming macro-clusters, which occurs with M1 macrophages. By contrast, slower and less persistent migration does not lead to significant clustering.

### Stiff matrices enhance migration speed but disable epithelial clustering for all macrophages

According to our results thus far, on soft matrices, epithelial cells migrate faster, more persistently, and form macro-clusters when co-cultured with M1 (but not M0 or M2) macrophages. To further explore this association between faster migration and clustering, we next asked whether stiff matrices could enable clustering because those are known to enhance epithelial migration due to mechanotransduction (2, 25). We fabricated stiff (120kPa) PA hydrogels and repeated experiments, as described above for soft gels. After time-lapse imaging for over 60hr on stiff matrices, we readily observed that epithelial cells did not form any clusters regardless of the macrophage type (Movie 4-6). The spatial and temporal distribution of nuclear pixel intensity remained unchanged overtime on stiff matrices (Figure 4A), indicating that epithelial cells remained as monolayers in contact with the matrix and did not cluster onto one another. Over the observed 60hr time-period, epithelial cells maintained their intercellular distance (Figure 4B), which also indicates that entire epithelial colonies move as a monolayer collective and do not undergo local clustering. In spatial distribution of cell velocity vectors, we noticed that cells near the leading edge were generally faster than those in the interior (Figure 4C), resembling conventional monolayer migration (2). Here, we found that cells seeded with M0-like macrophages travelled the farthest and maintained higher speeds at the front of the monolayer throughout the experiment (Figure 4A.i and 4A.ii). While epithelial cells co-cultured with M1 and M2 also started with higher cell speeds at the leading edge (Figure 4C.iii and Figure 4C.v, respectively), this speed advantage diminished over time (Figure 4C.iv and 4C.vi, respectively). Upon averaging migration parameters over time and samples, we found that epithelial cells co-cultured with M0 macrophages migrated with higher speed, persistence, and net displacement compared to M1 or M2 macrophages (Figure 4D, 4E, and 4F). Compared to soft matrices (Figure 4D, 4E, and 4F; dotted gray box), migration speeds were generally higher on stiff matrices, which could be too fast and not persistent enough to allow for cell-cell clustering. Thus, we argue that fast migration alone does not predict cell clustering. Instead, epithelial cells may require an optimal balance of migration, persistence, and cell-ECM adhesions to form stable clusters such that cells migrate enough to interact with one another but not break those intercellular connections due to prohibitively fast speeds.

**Figure 4.**
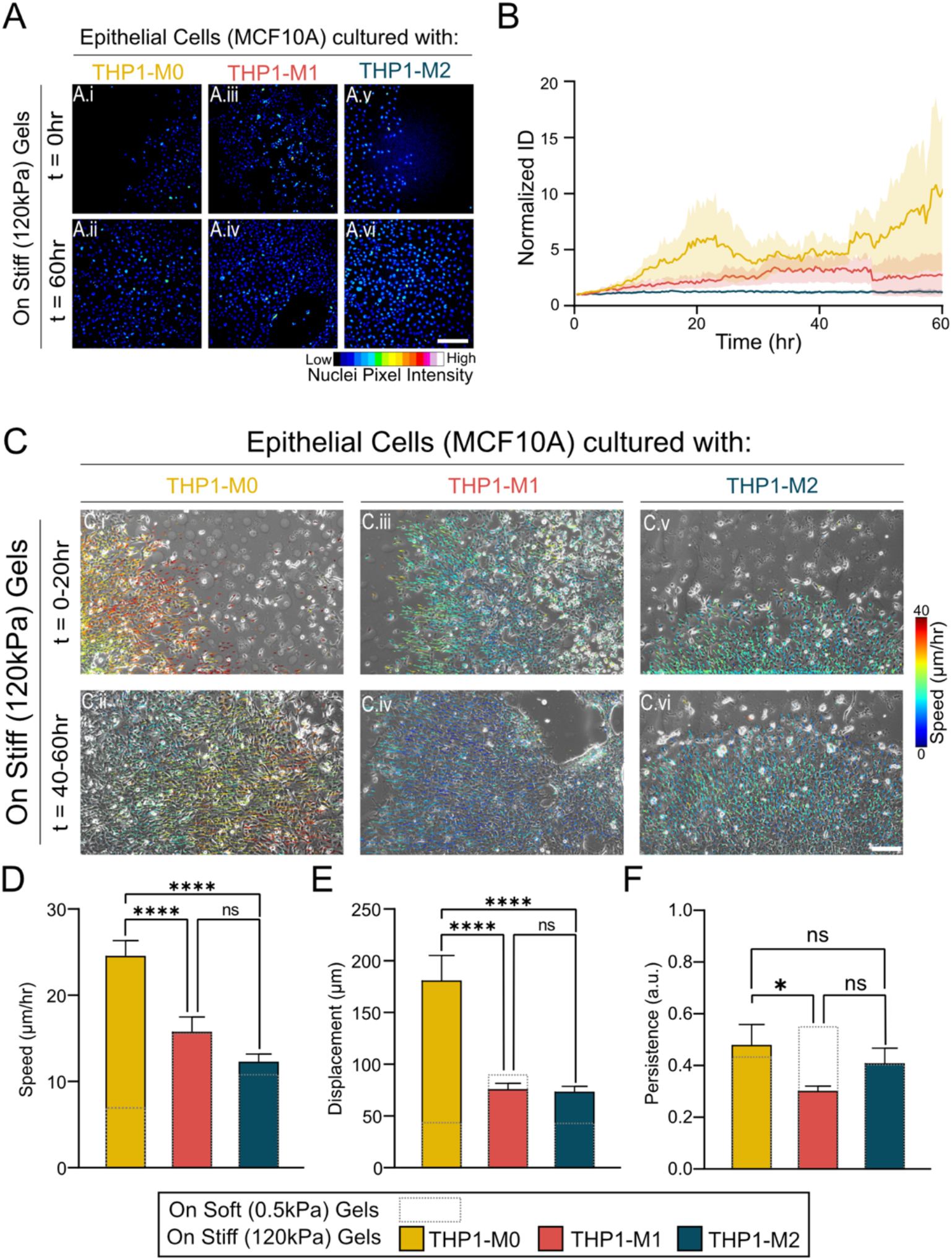
Stiff matrices disable macrophage-dependent epithelial clustering. **(A)** Representative images of MCF10A nuclei when co-cultured with (A.i and A.ii) M0, (A.iii and A.iv) M1, and (A.v and A.vi) M2-like macrophages on stiff matrices. **(B)** Line plot of normalized intercellular distance (ID) for MCF10As on stiff substrates in the co-presence of M0s, M1s, and M2s. Cells maintained their distance with other cells (measured as ID) over time regardless of macrophage type, indicating loss of clustering. **(C)** To understand migration and speed, quiver plots were overlayed with brightfield images of MCF10A cells between 0-20hr and 40-60hr time windows for (C.i, C.ii) M0, (C.iii, C.iv) M1, and (C.v, C.vi) M2-like macrophages. Quiver magnitudes depict cumulative distance over the 20hr window and colors depict average cell migration speed. Bar plots represent average **(D)** Speed, **(E)** Displacement, and **(F)** Persistence during peak clustering hours (t = 20-40hr), and error bars represent SEM. The overlayed dashed lines represent the soft counterparts (plotted earlier in Figures 2,3) during the same time window (t = 20-40hr). Data was analyzed by One-way ANOVA (with Tukey’s multiple comparison) for n≥3. ****P<0.0001, **P<0.01, *P<0.05, ns = not significant.

### Cluster formation facilitated by low cell-ECM adhesion formation and high fibronectin expression

During microtissue formation *in vitro*, different ratios of cell-ECM and cell-cell adhesions can cause tissue spreading versus cell aggregation (23). Substrate stiffness has also been shown to facilitate stronger cell-ECM adhesions, allowing for cells to spread and not aggregate (11). Here, to understand whether matrix stiffness and the type of co-cultured macrophages alter ECM adhesions of epithelial cells, we stained for paxillin, a strong protein marker in the focal adhesion (FA) complex (7), and measured the intensity minima and maxima to quantify number of separate adhesion bodies (see Methods and Figure 5A-C). On soft matrices, epithelial cells with M2 macrophages have significantly higher number of punctate focal adhesions, compared to those with M0 or M1 macrophages (Figure 5D.i-iv). In comparison, epithelial cells on stiff matrices had higher number of focal adhesions with no significant difference due to macrophage type (Figure 5E.i-iv). From these results, we infer that the condition that allowed formation of large epithelial clusters – M0 and M1 macrophages and soft matrix – has fewer focal adhesions, indicating that weaker cell-ECM interaction could benefit epithelial clustering. This is consistent with the classic differential adhesion hypothesis (26), which predicts higher cell-cell clustering and lowered spreading in cell populations with weaker cell-ECM adhesions.

**Figure 5.**
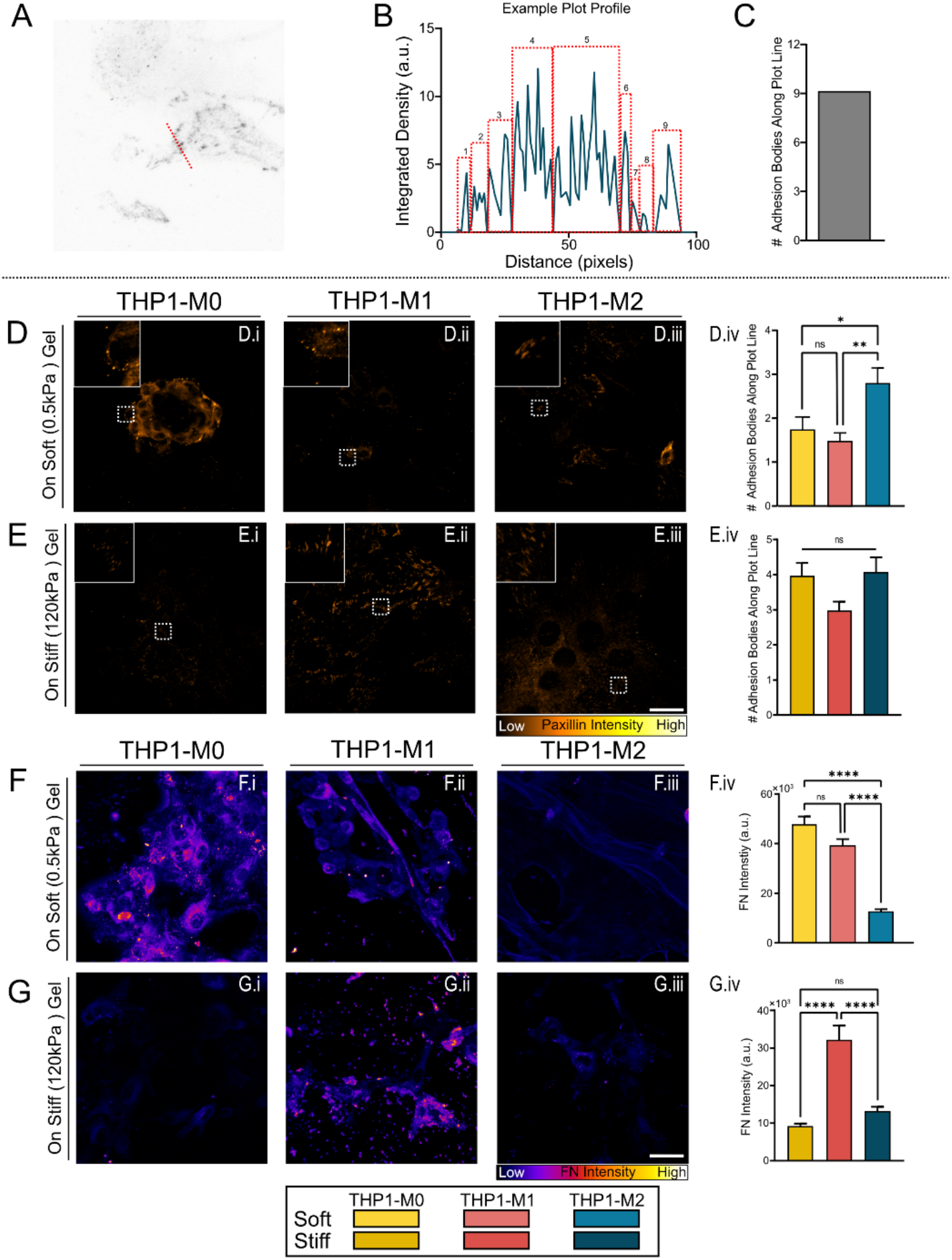
Adhesions and fibronectin in epithelial clusters. **(A)** To quantify stable paxillin punctate spots, we drew a line 100px in length. **(B)** Using image PlotProfile, we got a line plot of pixel intensity along the dashed line. When the intensity local minima hit 0, the section was counted as a paxillin punctate (red dashed rectangle). **(C)** The rectangles were then counted and plotted as the number of separate adhesion bodies along the plot line for n>25 cells in triplicate. Samples were fixed 48 hours after seeding epithelial cells. **(D)** Representative images of paxillin staining in co-cultures with (D.i) M0, (D.ii) M1, and (D.iii) M2-like macrophages on soft gels. Scale bar represents 20 *μ*m. Inset shows enlarged image of representative paxillin punctate spots. (D.iv) Bar plot illustrating the number of separate adhesion bodies measured along a plot line. Cells co-cultured with M2-like macrophages had the highest number of separate adhesion bodies compared to cells co-cultured with M0 or M1. **(E)** Representative images of paxillin staining of cells seeded on stiff (120kPa) PA gels. Cells were co-cultured with (E.i) M0, (E.ii) M1, or (E.iii) M2-like macrophages. Inset show zoomed in image of representative paxillin punctate spots. (E.iv) Bar plot representing the number of separate adhesion bodies along a plot line. On these stiff gels, there is no statistical difference in the number of separate adhesion bodies regardless of macrophage type. **(F)** Representative images of fibronectin (FN) staining of cells in (F.i) M0, (F.ii) M1, or (F.iii) M2 co-cultures on soft gels. Scale bar represents 20 *μ*m. (F.iv) Bar plot illustrating the average intensity of fibronectin expression. There was no statistical difference between the three soft conditions, regardless of macrophage type. **(G)** Representative images of fibronectin (FN) staining of cells in (G.i) M0, (G.ii) M1, or (G.iii) M2 co-cultures on stiff matrices after 48 hr. Scale bar: 20 *μ*m. (G.iv) Bar plot illustrating the average intensity of fibronectin expression. Data was analyzed by One-way ANOVA (with Tukey’s multiple comparison) for n≥2. ****P<0.0001, **P<0.01, *P<0.05, ns (not significant).

According to previous work, fibroblasts seeded on collagen type-I substrates with exogenously added fibronectin (FN) formed multicellular structures (14). Additionally, multicellular cluster formation via FN is due, in part, to weakened cell-ECM adhesions (15), consistent with our observations (Figure 5D.iv). Thus, we wondered whether fibronectin deposited by cells correlates with their clustering, depending on matrix stiffness and co-cultured macrophage polarization. We stained and quantified cellular FN expression. On soft gels, epithelial cells with M0 and M1 co-cultures had higher expression of FN compared to M2 co-cultures (Fig 5F.i-iv). On stiff gels, FN deposition was overall lower compared to soft matrices (Figure 5G.i-iv). On stiff gels, only epithelial cells with M1 co-cultures expressed high amounts of FN (Figure 5G.ii). Previously, fibronectin has been shown to aid in cell adhesion (27) and self-assembly of micro-tissues (14). On stiff gels, although M1 co-culture led to high FN deposition (Figure 5G.ii), it also associated with high cell-ECM adhesions (Figure 5E.ii) – a condition that likely does not assist in epithelial clustering. Taken together, these results suggest that macro-cluster formation on soft matrices with M1 macrophage co-culture is associated with relatively weaker cell-ECM adhesions and high FN deposition. When any one of those are out of balance, due to either macrophage type or matrix stiffness, the macro-cluster formation is disrupted.

### Myosin-IIA is highly expressed in cluster conditions, and ROCK inhibition abrogates clustering

Thus far, we have demonstrated two distinct stiffness- and macrophage-dependent cluster types. Instantaneous speed and persistence are higher during macro-clustering than in micro-clustering on soft matrix. Moreover, cell clustering on soft substrates requires lower cell-ECM adhesions and higher FN expression. However, it is not clear whether intracellular contractility contributes to stiffness-dependent cluster formation. Previous work has attributed formation of multicellular structures to RhoA- and actomyosin-dependent cell contractility (11, 22). Therefore, we measured expression of non-muscle myosin IIA (NM2A) and alignment of filamentous actin (F-actin) fibers. After seeding MCF10As in the presence of macrophages on soft gels for 48hr, cells were fixed and stained for NM2A and F-actin (Figure 6A). To quantify NM2A expression across samples, we normalized NM2A signal to Hoechst and performed measurements within ROIs of 500*μ*m^2^ area throughout the clusters. On soft matrices, NM2A intensity in epithelial cells was significantly higher when cultured with M1 macrophages, compared to M0 and M2 (Figure 6B). In addition, F-actin intensity was highest on soft matrices in cells seeded with M0s or M1s (Figure 6C). Although actin intensity was higher in cell clustering conditions, there was no significant difference in actin fiber coherency (a measure of actin fiber alignment) in the soft conditions (Figure 6D). During micro-cluster formation, epithelial cells cultured with polarized macrophages (M1 or M2) generated lower traction forces compared to those with M0 (Figure 6E), further indicating reduced interaction with the ECM.

**Figure 6.**
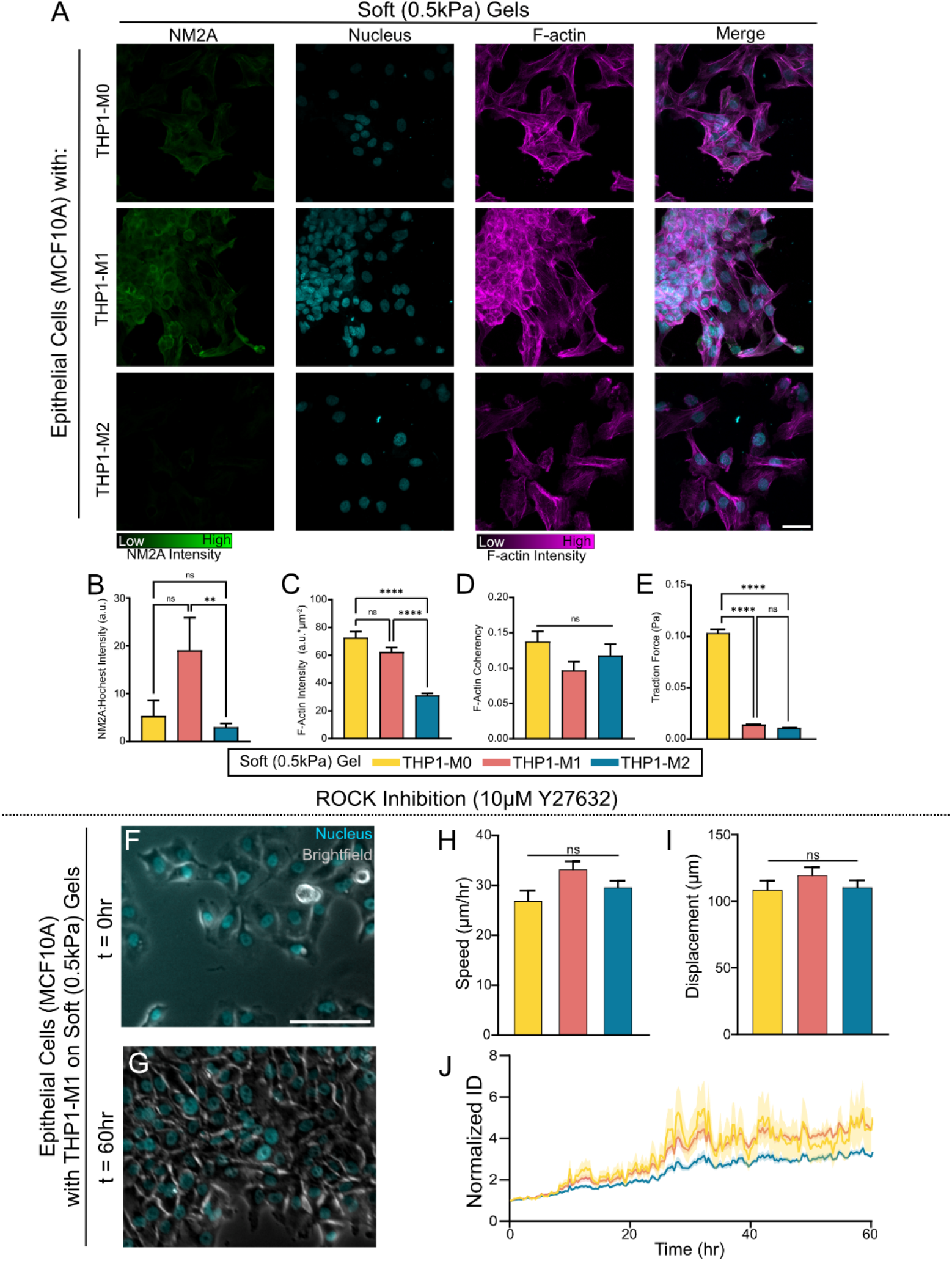
Cellular contractility required for cluster formation. **(A)** Representative images of non-muscle myosin IIA (NM2A), nuclei, F-actin, and merged images of epithelial cells co-cultured with different activated macrophages on soft gels. Color scales represent pixel intensity, scale bar represents 20µm. Representative images show the highest NM2A intensity in epithelial cells co-cultured with M1-like macrophages on soft gels – the condition that enabled macro-clustering. **(B)** Pixel intensity of NM2A (normalized to nuclear intensity to account for any changes in imaging across samples) measured in epithelial clusters (n > 8 clusters). **(C)** Actin intensity measured in cell clusters, normalized by ROI area (n > 30 cells). **(D)** Using OrientationJ, ImageJ plugin, coherency of F-actin fibers (a measure of fiber alignment) was calculated in at least two different regions of n = 30 cells. Actin coherency was similar for all soft conditions regardless of macrophage type. **(E)** Tractions exerted by the cells were measured using Traction Force Microscopy (TFM). To test whether loss of contractility affects clustering, ROCK expression was chemically inhibited using 10µM Y27632. Representative images of epithelial cells (nucleus in cyan) in the presence of M1-like macrophages on soft gels at **(F)** t = 0hr and **(G)** t = 60hr. Unlike the control (untreated) samples, the epithelial cells did not form clusters by 60hr. Scale bar represents 100µm. Bar plots of average **(H)** speed and **(I)** displacement show ROCK inhibitor causes cell migration to not vary with macrophage type. Error bars=SEM. **(J)** Normalized intercellular distance (ID) steadily rose regardless of macrophage type. Chemical inhibition of ROCK allowed epithelial cells to expand outwards and not form clusters. Line represents average values and shaded regions represent SEM. Data was analyzed by One-way ANOVA (with Tukey’s multiple comparison) for n = 3, unless otherwise stated. ****P<0.0001, **P<0.01, *P<0.05, ns (not significant).

Consistent with previous work, our findings indicate that intracellular contractility relates to cell aggregation and cluster formation (12, 28, 29). Comparatively, on stiff matrices, there were little differences in NM2A and F-actin expressions across macrophage types (Figure S4B-D). To further understand the role of contractility in cluster formation, we performed pharmacological inhibition of Rho-associated kinase (ROCK) using 10*μ*M Y27632 after cell seeding. Indeed, as hypothesized, epithelial clusters did not form after ROCK inhibition (Figure 6F-G) on soft matrices, which showed macro- or micro-clustering without treatment. Instantaneous migratory speed (Figure 6H) and net displacement (Figure 6I) of ROCK-inhibited epithelial cells did not change with macrophage type. We also observed that cells increasingly move away from one another, quantified by intercellular distance >1 for the duration of experiment (Figure 6J). Overall, these results show that myosin expression (regulator of cellular contractility) correlates with epithelial clustering and loss of ROCK activity (required for active mechanotransduction) disables both micro- and macro-clustering.

### Cytokine secretion correlates to epithelial clustering on soft matrices

While some previous studies have explored self-assembly of microtissues and cellular aggregate formation using cell culture *in vitro* (15, 22, 23), our work also accounts for co-presence of macrophages with cues from soft substrates. An important known function of macrophages is cytokine secretion to facilitate wound healing, chemotaxis of other immune cells, tissue remodeling, and angiogenesis (30). The types of cytokines released is dependent on the type of macrophage, such as TNF-α from M1-like macrophages and TGF-β from M2-like macrophages (21, 30). We wondered whether the observed differences in epithelial clustering could be attributed to different cytokine secretion corresponding to macrophage types. We measured cytokine concentration for TNF-α and TGF-β using sandwich enzyme-linked assay (ELISA). We chose both cytokines not only because they are secreted by macrophages but also because TNF-α has been linked to collagenase receptor inhibition (31) and TGF-β has been linked to pro-tumor growth (32). We measured cytokine concentration in supernatant collected from co-cultured epithelial cells with polarized macrophages on PA gels after 72hr. We found that TNF-α was secreted on soft matrices with all three types of macrophages, with highest levels in the presence of M1 macrophages (Figure 7A) – condition that supported macro-clustering. We also found that TGF-β secretion was only detected in MCF10A cells with M2 co-culture on soft substrates (Figure 7B); however, the co-cultures with M0 and M1 macrophages were below the limit of detection (<10pg/mL).

**Figure 7.**
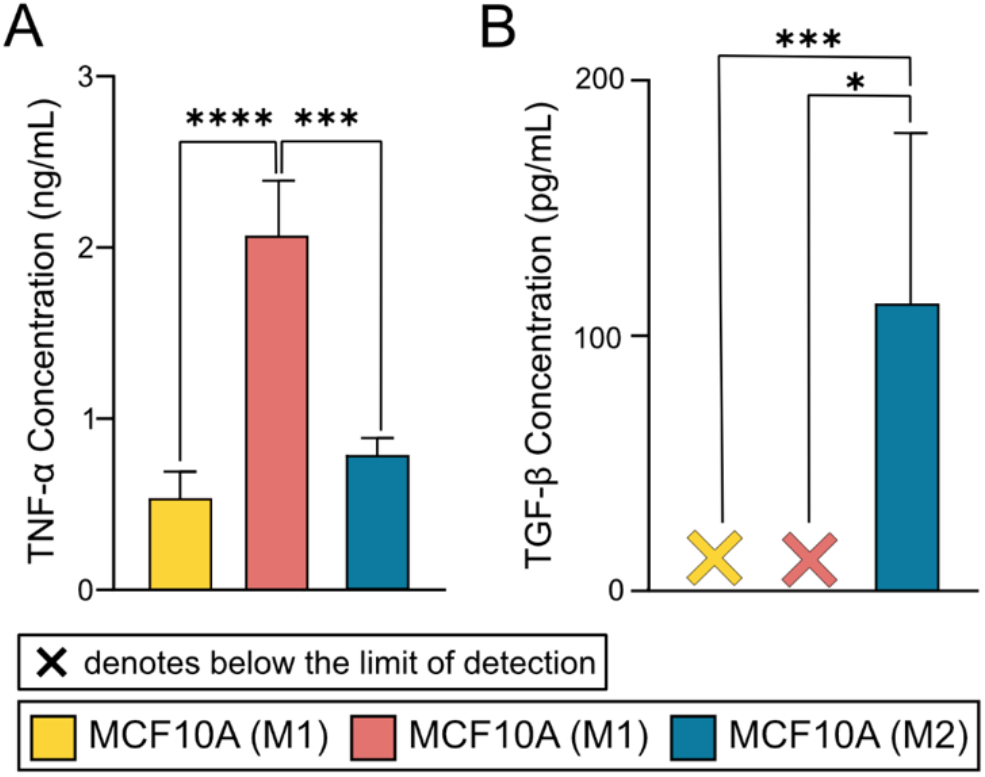
Cytokine secretion measurements from ELISA on soft substrates. Cytokine secretion of **(A)** TNF-α and **(B)** TGF-β in co-cultures of MCF10A cells with macrophages of M0, M1, and M2 polarities. “X” denotes measurements were below the limit of sensitivity (<10pg/mL). Data was analyzed with One-way ANOVA (with Tukey’s multiple comparison) from n=6 ran in duplicate. ****P<0.0001, ***P<0.001, *P<0.05.

### Addition of TGF-β accentuates epithelial clustering

After measuring cytokine secretion, we wondered how the addition of cytokines would impact epithelial clustering with macrophage co-culture. Previous work has shown TNF-*α* and TGF-β can synergistically promote EMT (32). To test this hypothesis, we added 25ng/mL TGF-β in separate experiments. Upon the addition of exogenous TGF-β, we observed that micro-clusters formed on soft matrices in MCF10A epithelial cells in the co-presence of M0 macrophages (Figure 8A). These micro-clusters stayed intact over the 60hr duration of live imaging (Figure 8A). In the co-presence of M1 macrophages, MCF10A cells started as spread out, but eventually formed macro-clusters by the end of the 60hr experiment duration (Figure 8A). Cells in co-culture with M2 macrophages underwent some clustering; however, these clusters contained fewer cells compared to micro-clustering observed in case of co-cultures with M0s on soft gels (Figure 8A). Clustering is further confirmed in measurements of Normalized ID, where conditions maintain distance at or below one (Figure 8B), indicating that cells came closer overtime. Similar to the control experiments (without TGF-β), macro-cluster formation was associated with both higher instantaneous speed and net displacement compared to cells forming micro-clusters (Figure 8C-D). Thus, we argue exogenous addition of TGF-β amplifies clustering, specifically macro-clustering, because of the decrease in Normalized ID overtime, and higher migratory speed and displacement.

**Figure 8.**
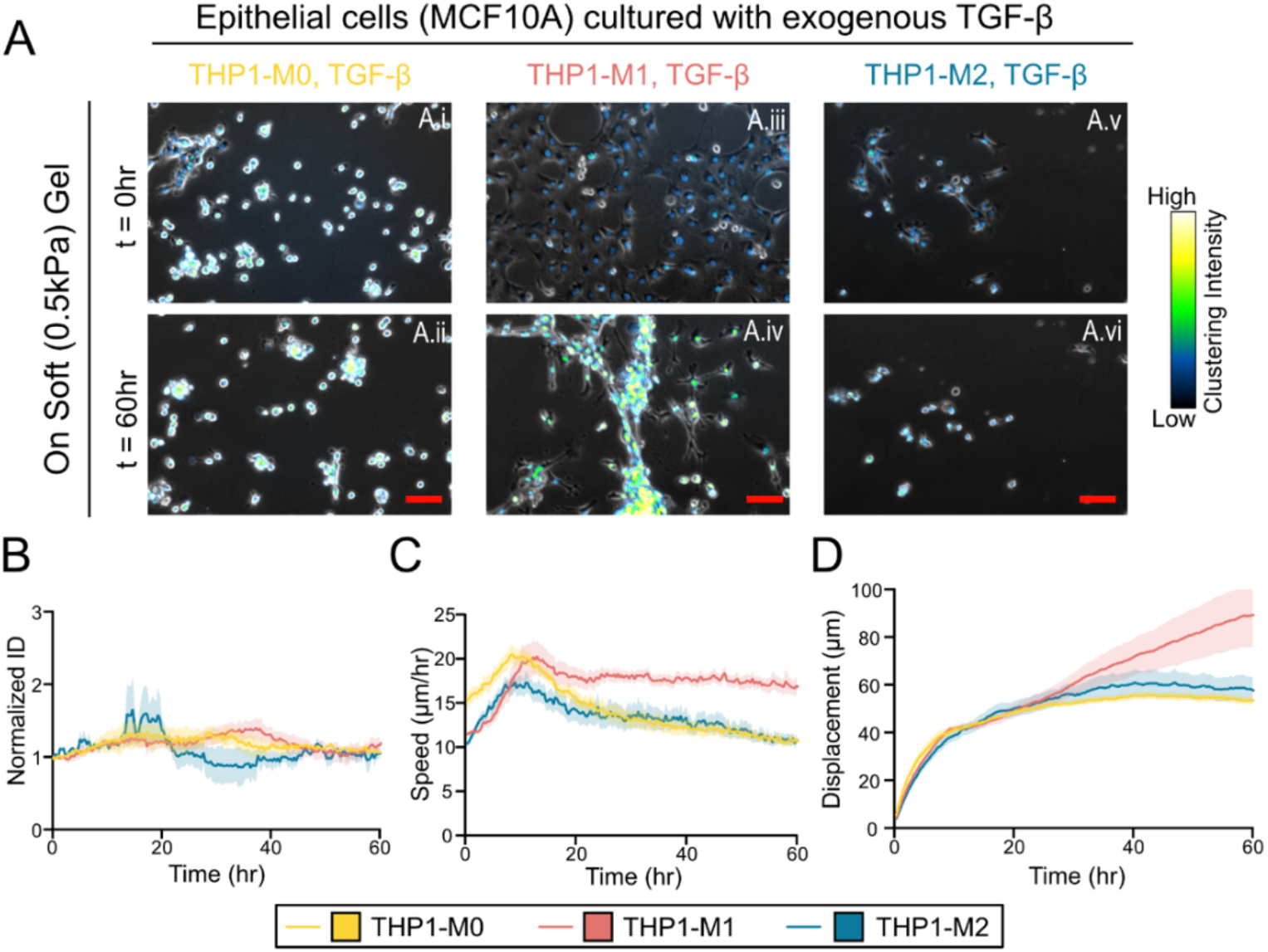
Epithelial clustering after exogenous addition of TGF-β. **(A)** Representative images of cells in the presence of 25ng/mL exogenous TGF-β on soft matrices. Brightfield images merged with epithelial cell nuclei with pixel intensity at t = 0 hr for (A.i) M0 at t = 0hr and (A.ii) t = 60hr; (A.iii) M1 at t = 0hr and (A.iv) at t = 60hr; and (A.v) M2 at t = 0hr and (A.vi) t = 60 hr. Scale bar represents 100µm and color bar represents epithelial nuclei pixel intensity. Line plots for **(B)** Normalized ID, **(C)** Speed, and **(D)** net displacement overtime in the presence of exogenous TGF-β. Line plots represent average and shaded region represents SEM for n = 3.

## Discussion

Cellular cluster formation has previously been studied from both biochemical and mechanical perspectives (6, 11, 12, 23, 33). It has been shown that multicellular clusters form on softer substrates but not on stiff substrates (11). Although heterogeneous tissue environments include immune cells that dynamically interact with epithelial cells, it is less understand whether immune cells can affect cell clustering. In this study, we show for the first time that epithelial cluster formation is regulate by macrophage polarization and substrate stiffness (Figure 9). We found that soft substrates in combination with M1-like macrophages cause macro-cluster formation, while M0 macrophages promote micro-clusters. In conditions that support cluster formation, cell-ECM adhesions were weak and fibronectin expression was high. Furthermore, formation of large clusters (macro-clusters) in the presence of M1 macrophages on soft substrates was associated with higher actin-myosin expression and TNF-α secretion (Figure 9). By contrast, stiff substrates disabled cluster formation because cell-ECM adhesions could be too strong to break and migrate speeds were too fast to enable stable cell-cell interactions (Figure 9). These results illustrate how the co-presence of macrophages can tap into conventional mechanosensing processes (adhesions, contractility) to interrupt normal epithelial migration on soft matrices and promote clustering.

**Figure 9.**
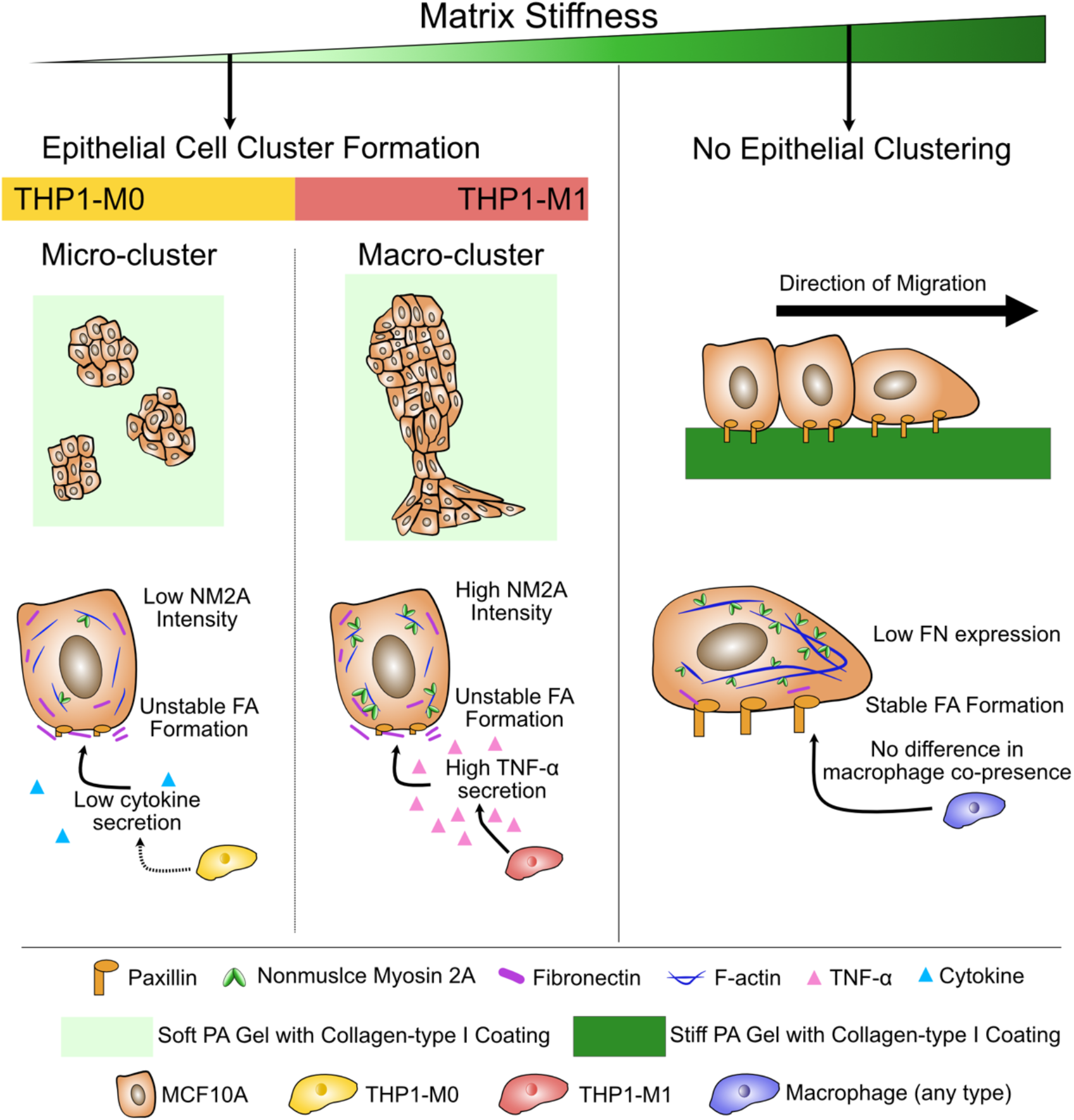
Summary of regulators and mediators of epithelial clustering by macrophage polarity and matrix stiffness. Epithelial cluster formation reduces for stiffer matrices. <u>Left panel</u>: On soft matrices, epithelial clusters form but their geometry varies with types of co-cultured macrophages. With M0-like macrophages, epithelial cells form small ‘micro-clusters’, which are smaller in area and shape index, express lower NM2A, make stable focal adhesions (FAs), and secrete lower levels of cytokines. Alternatively, with M1-like macrophages, epithelial cells form macro-clusters (larger in area and shape index), express higher NM2A and FN expression, make unstable FAs, and secrete higher levels of TNF-α cytokine. Upon ROCK inhibition on soft matrices, cluster formation is abrogated due to loss of contractility. <u>Right panel</u>: On stiff matrices with all three types of macrophages, epithelial cells maintain sheet-like monolayers, make stable FAs, express lower levels of FN, and migrate faster, all of which contributes to loss of multicellular clustering.

During organ development, as tissues begin to assemble, cells of similar physical properties must come together, which has been described using a Differential Adhesion Hypothesis wherein cohesive cells cluster together while cells with stronger ECM adhesions move away, analogous to mixtures of immiscible liquids (34). Alternatively, the Differential Surface Contraction Hypothesis argues that inherent differences in actomyosin-dependent tension regulate cell aggregate formation (35). Cellular cluster formation has also been described using physical principles of active wetting of fluids (12, 26, 29), influenced by substrate stiffness (11, 12, 29). In this study, we note that cell cluster formation is not a static process; rather, cells dynamically move and explore their environment prior to clustering (12, 26). According to our findings, these pre-clustering steps of migration, cell-ECM adhesions, and cell-cell interactions depend on matrix stiffness and co-cultured macrophages. We found that soft matrices and M1 macrophages generate an optimal balance of weak cell-ECM adhesions and slower migration while maintaining actin-myosin contractility. When any of these components are disrupted, cell clustering is diminished or eliminated. According to previous work, traction forces generated at the periphery of multicellular clusters decrease overtime with the cluster size (12), indicating that cellular forces are necessary for stable clustering. In our study, both the loss in cellular contractility by ROCK inhibition (Figure 6J) and the activation of cell migration on stiff matrices (Figure 4C) disable clustering.

Additional factors, including substrate stiffness (11, 12), extracellular matrix density (28), cluster size (12), and cluster density (12) can all affect tissue wetting as well as the cell dynamics and forces. In this study, we found two different cluster types: *“micro-clustering”* and *“macro-clustering”* Both cluster types require weak cell-ECM adhesions, as seen in the low number of separate paxillin punctate spots (Figure 5D.iv). On stiff matrices, stable focal adhesions of cells with the matrix (Figure 5D.iv) possibly work against cell-cell cohesion and clustering. Overall, our findings show that epithelial clustering is driven by weak cell-ECM adhesions and high contractility (Figure 9). While the influences of substrate stiffness on cell-cell cohesions and scattering have previously been studied (29), our findings reveal co-culture macrophage type can provide a new crosstalk with mechanosensing and effect epithelial clustering. This contribution from macrophages could be from their cytokine secretion, like TNF-α measured in Figure 7A. In future studies of disease (cancer, fibrosis) or wound healing, where epithelial cell clustering is crucial and cell reside in complex mechanical and immune environment, contributions from polarized macrophages in concert with matrix stiffness could be considered. This additional lever of macrophage polarity could provide novel ways to tune biophysical evolution of multicellular clusters.

## Methods

### Polyacrylamide Gel Synthesis and Collagen Type I Coating

Soft and stiff polyacrylamide gels were fabricated on glass bottom well plates for live time-lapse microscopy and glass coverslips for fixed imaging. Polyacrylamide precursor solutions were mixed by adjusting 40% acrylamide (Bio-Rad: 161-0142) and 2% bis-acrylamide (Bio-Rad: 161-0140). Soft (0.5kPa) and stiff (120kPa) PA gels were made by mixing 4%/0.2% and 15%/1.2%, respectively (25, 36). Gels were polymerized on glass bottom plates for live time-lapse microscopy and glass coverslips for fixed immunofluorescence experiments. Glass surfaces were activated by plasma coating for 10 minutes then adding Bind Silane solution (95% EtOH, 5% Acetic Acid, 0.3% Bind Silane; GE Healthcare: GE17-1330-01) for 10 minutes. Glass surfaces are then sprayed with 70% ethanol and dried using a Kimwipe.

To make PA gels in glass bottom plates, 5% Ammonium Persulfate (Sigma-Aldrich: 161-0700) and 0.5% Tetramethylethylenediamine (TEMED; Sigma-Aldrich: T7024) were added to 1mL PA gel solution and gently mixed. Once mixed, 20μL of PA gel solution was added to the glass surface and immediately covered with Sigmacote (Sigma-Aldrich: SL2) treated glass coverslips. Gels were allowed to polymerize for at least 30min before adding PBS. To make coverslip gels, 24-28μL gel solution was added to Sigmacote-treated glass slides and glass coverslips were immediately added on top of the PA gel solution. Coverslips were immersed in PBS until used. PA gel surfaces were functionalized using sulfo-SANPAH (Thermo Fisher: 22589) and activated under UV light for 10min. Plates were washed twice with sterile PBS before adding 0.05mg/mL Collagen type-I (Santa Cruz: SC-136157). Plates were covered and left in the 4°C overnight. Plates were washed with sterile PBS and air-dried in the hood for 10 minutes before seeding cells.

### Cell Culture

Human monocyte cell line (THP-1) (gifted from Dr. Azab laboratory; University of Texas Southwestern Medical Center, Dallas, Texas) was cultured in RPMI-1640 (R8758; Sigma, St. Louis, MO, USA) with 10% fetal bovine serum (FBS; Thermo Fisher: MT35016CV) and 0.2% Normocin (Thermo Fisher: NC9273499). Monocytes were not used until the third passage and not used past the seventh passage. Human mammary epithelial cell line (MCF10A) was cultured in DMEM/F12 (Cytivia: SH30023.01) with 5% horse serum (Sigma Aldrich: H1138-500ML), 0.2% Normocin, 20ng/mL epidermal growth factor (EGF; Miltenyi Biotech: 130-097-750), 0.5mg/mL hydrocortisone (Sigma-Aldrich: H0888), 100ng/mL cholera toxin (Sigma-Aldrich: C8052), and 10*μ*m/mL insulin (Sigma-Aldrich: I6634). Cells were not used until past three passages.

Monocytes were activated to macrophages by adding 30ng/mL phorbol-12-myristate-3-acetate (PMA; Sigma-Aldrich: P8139) for 24hr. Macrophage polarization varied based on macrophage type and was based previous work (37). Naïve macrophages (M0) were rinsed and allowed to sit in THP-1 growth media for 48 hr. M1-like macrophages were activated by adding 5ng/mL interferon gamma (IFN-*γ*; PeproTech: AF-300-02) and 10ng/mL lipopolysaccharide (LPS; Sigma-Aldrich: L2630) for 48hr. M2-like macrophages were activated by adding 25ng/mL interleukin 4 (IL-4; PeproTech: 200-04-50UG) and 25ng/mL IL-13 (PeproTech: 200-13-50UG). Once activated, macrophages were rinsed with warmed sterile PBS before adding 18,000 cells were seeded on PA gels. The MCF10A monolayer was made based on the previously used bubble method (38). Epithelial cells were allowed to adhere for 25min atop macrophages. After adding media, MCF10As were allowed to acclimate for 6hr before live time-lapse experiments or 48hr before fixed experiments.

### Live Timelapse Microscopy and Quantitative Analysis

Following 6hr in cell culture, live time-lapse microscopy experiments were started on an Olympus microscope (Olympus, Evident, Tokyo, Japan). Phase contrast and green fluorescence images were acquired at 20min intervals for 60-72hr at 10x objective. Stage incubator was maintained at 37°C and 5% CO2 during the experiment. After imaging, experiment tiffs were exported from the microscope. To reduce bias, experiments were repeated at least three times with at least five positions taken within each sample condition.

Image stacks were analyzed using Fiji ImageJ (NIH, Bethesda, MD, USA). Cluster analysis was done by taking final frame of a stack and manually tracing in ImageJ. Cluster parameters were measured in ImageJ. Nuclear intensity of clusters was measured in 200×200px^2^ regions and average intensity measured within regions of interest. Cell migration analysis was performed by analyzing epithelial nuclei using ImageJ manual tracking during cluster events or plugin TrackMate (NIH, Bethesda, MD, USA) (39, 40). Raw data was processed through a custom Python code (41) using the following equations:

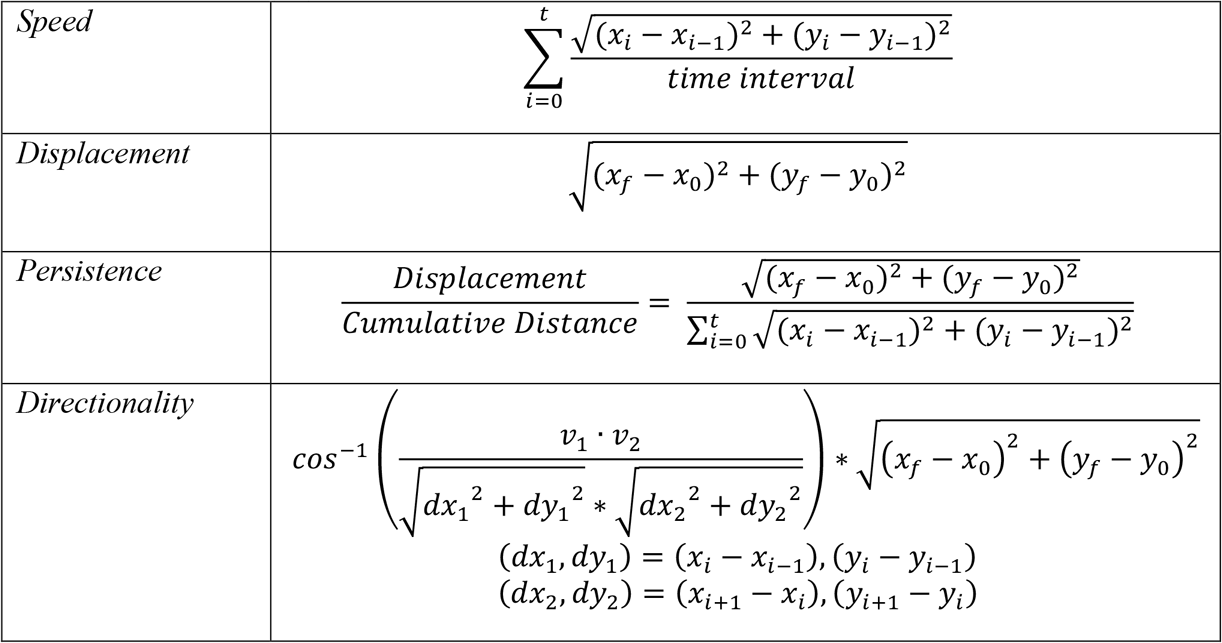

In addition to TrackMate and manual tracking, distance between the nuclei of the cell were tracked and quantified into Normalized Intercellular Distance (ID). ROIs of 250×250px^2^ were taken for each condition, at least three per image to reduce bias. Cell nuclei were tracked using StarDist2D (42) (Figure S1A). The distance was calculated using the following equation:

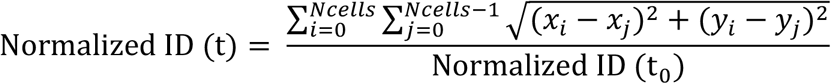

### Fixed Imaging and Immunofluorescence Analysis

For fixed images, cells were added to coverslip gels using the same method as in glass well plates. Cells were left in culture for 48hr before fixing. Cells were fixed by first washing the gels of media with warm sterile PBS once and then sat in 1mL 4% PFA for 10 minutes. Gels were then rinsed twice with Sterile PBS. Cells were then rinsed three times with 0.5% bovine serum albumin (BSA; Millipore Sigma: 126593) before permeabilization in 0.1% Triton X-100 (Sigma-Aldrich: 234729) in PBS for 10 min. The samples were then blocked with 1% BSA, 10% Goat Serum (Sigma-Aldrich: G29023-10ML) and 0.3M Glycine (Sigma-Aldrich: G8898-500G) for 1hr at room temperature. Primary antibodies were added based on the following dilutions: Anti-paxillin (1:200; Cell Signaling Technology: D7S2M), Anti-Fibronectin (1:400; Sigma-Aldrich: F3648-100UL), Anti-Catenin-δ1 (P120; 1:800; Cell Signaling Technology: 59854), and non-muscle myosin-2A conjugated with Alexa 647 (NM2A; 1:100; Abcam: ab204676-100UL).

Primary antibodies were diluted with 0.5% BSA in PBS and left in the dark at 4°C overnight. Afterwards, the plate was rinsed with 0.5% BSA in PBS twice before adding secondary antibodies. Secondary antibodies were diluted to 1:100 in PBS and left in the plate for 2hr at room temp without light exposure. The plate was then rinsed twice with PBS before adding Rhodamine Phalloidin (1:500; Invitrogen: R415) for 45min at room temp without light exposure. The plate was rinsed once with PBS and then Hoechst 33258 (20µg/mL) was added for 30 min at room temperature away from light. Samples were gently washed twice with PBS before mounting samples onto glass slides. Samples were mounted by covering the sample with a coverslip and sealing with transparent nonfluorescent nail polish. To prevent bias, at least two replicates were used with at least six positions taken per condition. Images were taken on a Zeiss LSM confocal microscope (Carl Zeiss MicroImaging, Oberkochen, Germany) with either 40x (oil objective) or 63x (oil objective). All samples had F-actin and Hochest used in the samples to label the cell body and nucleus.

#### Focal Adhesion Analysis

Focal adhesions were identified using the protein paxillin. Paxillin punctuates were analyzed in ImageJ. Raw z-stacks were projected along max intensity. To measure the number of separate adhesion bodies, a uniform plot line (∼100µm in length) was drawn across paxillin spots at the edge of the cell (Figure 5A). The local maxima peaks were counted, resulting the number of separate adhesions bodies along a plot line for the cell (Figure 5B-C).

#### Intensity Analysis

Maximum intensity was projected for each z-stack. At least three ROIs of 500*μ*m^2^ were chosen randomly and analyzed for Raw Integrated Intensity. Raw Integrated Intensity was either normalized by the average Hoechst intensity or ROI area.

#### Coherency Analysis

Coherency of F-actin fibers was analyzed using OrientationJ (43) ImageJ plugin by measuring the degree to which fibers aligned relative to other fibers in the ROI. At least three regions of 500*μ*m^2^ were randomly chosen.

### Traction Force Microscopy

To detect displacement field within the matrix, PA gels were made using the same protocol described above, but with the addition of 2μL of cy3 fluorescent beads (Thermo Fisher: F8820) to every 1mL of PA gel solution. Gels were flipped during polymerization to ensure that beads settle near the surface over which cells are seeded. Live cell imaging was performed after 72hr cell culture on gels, as described above. Images were taken of z-stacks covering 100*μ*m depth at 5*μ*m slice intervals. After initial image collection, 1mL 0.25% trypsin (Sigma-Aldrich: T4049) was added to each well and immediately reimaged.

Image stacks were analyzed by projecting maximum pixel intensity of beads across the z-axis. Images were aligned along the x-y axis using the Slice Alignment plugin in ImageJ (Template Matching and Slice Alignment, ImageJ). Lastly, bead displacement was measured using PIV Cross-correlation (PIV, ImageJ) plugin followed by Traction Force plugin (Traction Force Microscopy, ImageJ). Traction force magnitude was outputted and plotted in GraphPad Prism.

### Chemical Inhibitor and Cytokine Secretion

To inhibit, Rho kinase (ROCK) activity, 10µM Y27632 (Millipore Sigma: 688000-1MG) was added 6hr after seeding epithelial cells, followed by live timelapse imaging. Proliferation was inhibited using 1µM Itraconazole (Sigma-Aldrich: PHR1834-200MG) diluted in DMSO to 1mM and PBS to 1µM, added 24hr after cell seeding.

For ELISA, cell media was collected from plates 72hr after seeding epithelial cells. Supernatant was collected from at least six biological replicates according to manufacturer protocol. ELISA was run for TNF-α (Millipore Sigma: RAWB0476) and TGF-β (Millipore Sigma: RAB0460).

### Statistical analysis

All live experiments have at least three biological replicates and fixed images have at least three biological replicates, unless otherwise stated. Raw data was compiled in GraphPad Prism (GraphPad Software LLC, La Jolla, CA, USA). For statistical analysis, ANOVA (Tukey’s multiple comparison test) was performed to assess statistical significance between groups. Standard values for reporting significance were used. Data was plotted as mean and standard error of the mean (SEM), unless stated otherwise.

## Abbreviations

PA: polyacrylamide
ID: Intercellular Distance
ECM: Extracellular Matrix
FA: Focal Adhesion
NM2A: Non-muscle myosin
IIA; TNF-α: Tumor Necrosis Factor α
TGF-β: Tumor Growth Factor β
IL-4: Interleukin 4
IL-13: Interleukin 13
IFN-γ: Interferon γ
ELISA: enzyme-linked immunoassay

## Acknowledgments

This work was supported by the NIH/NIGMS MIRA (R35GM128764) grant to AP. H.Z. was supported by NSF GRFP (DGE-2139839 and DGE-1745038).

## Author Contributions

HZ and AP conceived the project and designed experiments. HZ performed experiments, analyzed data, and wrote the manuscript. HZ and AP designed experiments and interpreted results. AP edited the manuscript, supervised project, and acquired funding.

## Data Availability Statement

The data that support the findings of this study are available in the methods and/or supplementary material of this article.

