## Supplementary Figures for "Epithelial multicellular clustering enabled by polarized macrophages on soft matrices"

### Supplemental Figures

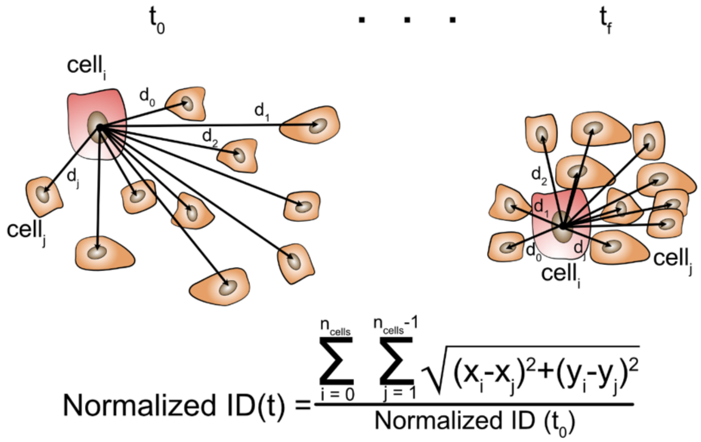


**Figure S1. Normalized Intercellular Distance (Normalized ID) measures degree of clustering.** Schematic of Normalized ID measurements from $t_0$ to $t_f$. As cells become clustered, the cell nuclei come closer together and produce Normalized ID below 1.

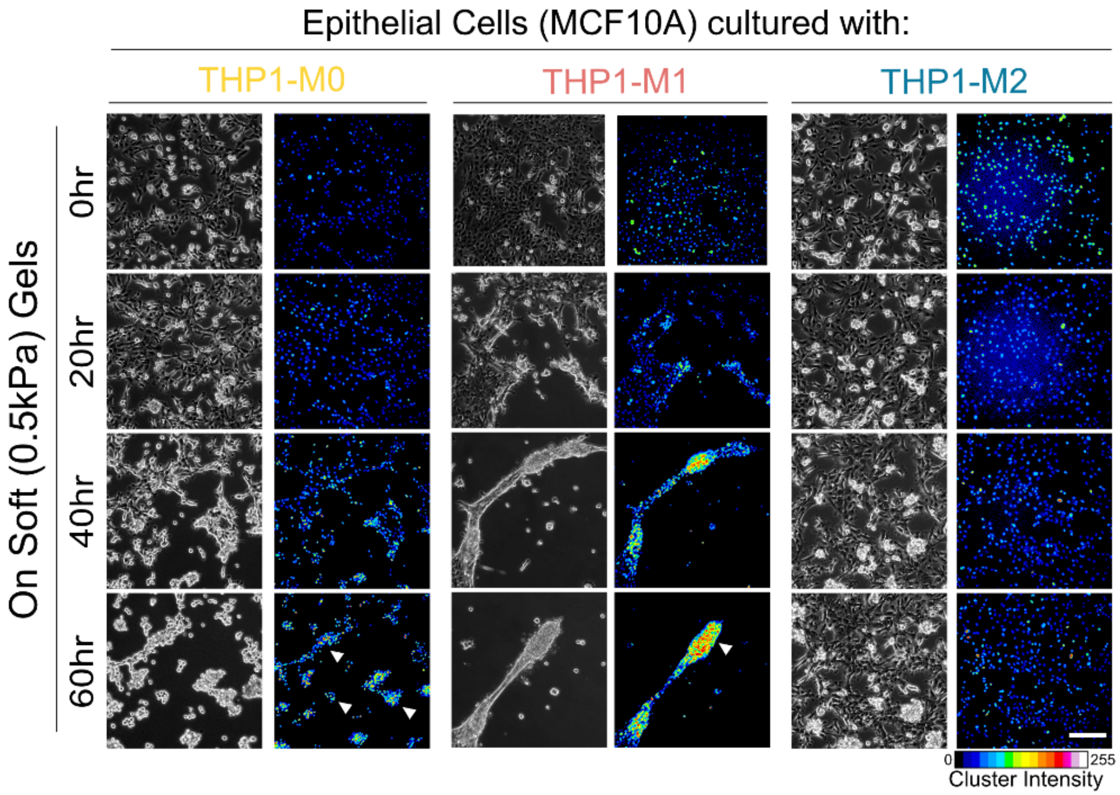


**Figure S2. Epithelial cell clustering over time on soft gels.** Representative images of epithelial cells in brightfield and fluorescing nuclei over the experiment time in the presence of M0, M1 or M2 macrophages, with largest clustering with M1 macrophages over the t = 60hr experiment time.

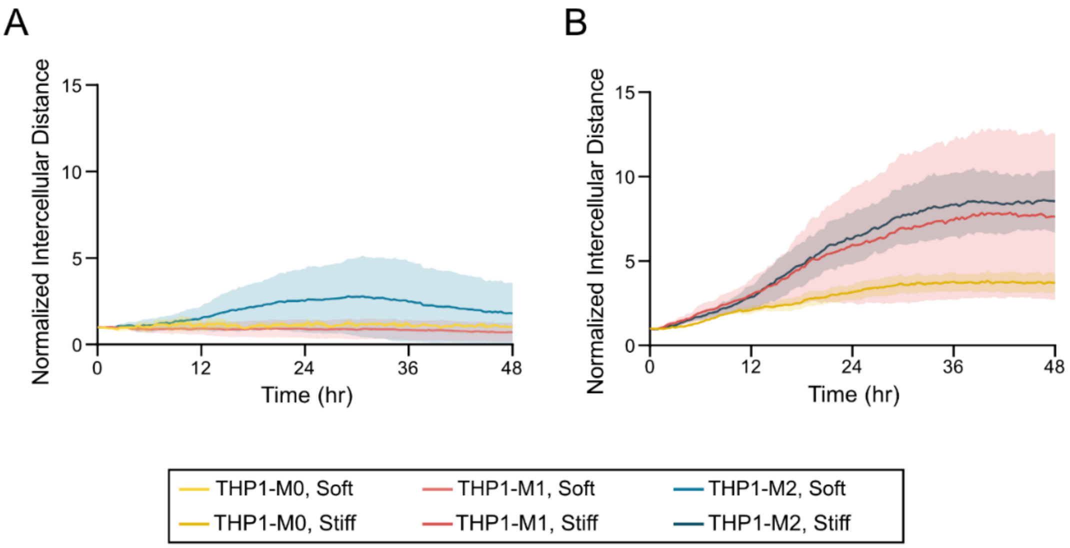


**Figure S3. Proliferation inhibitor preserves clustering phenotype.** Proliferation inhibitor, Itraconazole, was added to **(A)** soft and **(B)** stiff conditions. Line plots show there is little difference in Normalized ID from control setting. Line plot represents average and shaded region represents SEM for n=3.

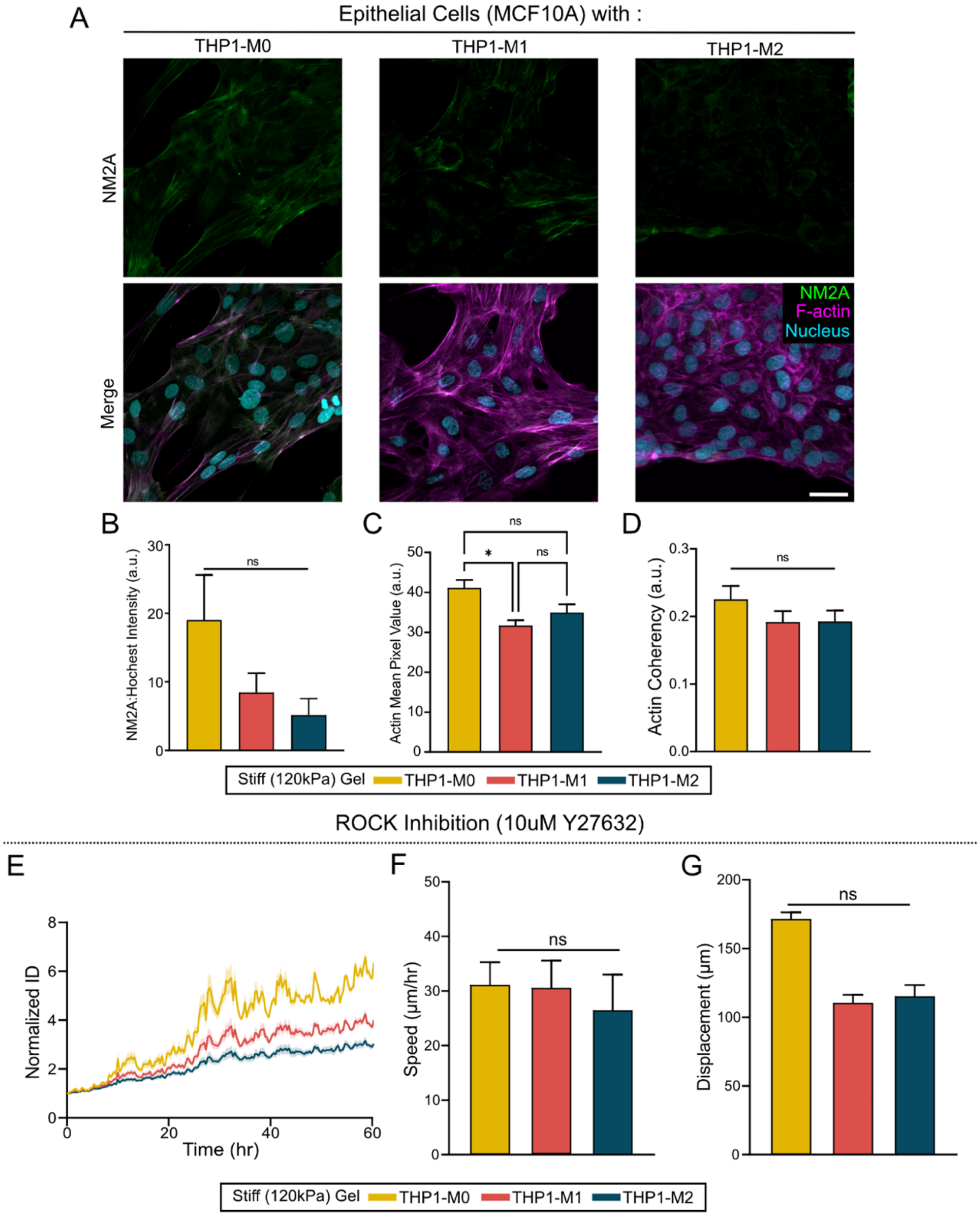


**Figure S4. Role of cellular contractility in epithelial behavior on stiff substrate. (A)** Representative fluorescence images of NM2A and merged channel with NM2A, nuclei, and F-actin. **(B)** NM2A intensity measurements normalized against Hoechst signal. (n > 25). **(C)** F-actin expression and **(D)** actin coherency in the co-presence of macrophages (n > 25). **(E)** Line plot representing the Normalized ID of epithelial cells co-cultured with macrophages and treated with 10µM Y27632 to chemically inhibit ROCK. Representative bar plots for **(F)** speed and **(G)** net displacement show no difference in migration of epithelial cells in the presence of difference macrophages on stiff matrices. *P<0.05, ns = not significant.

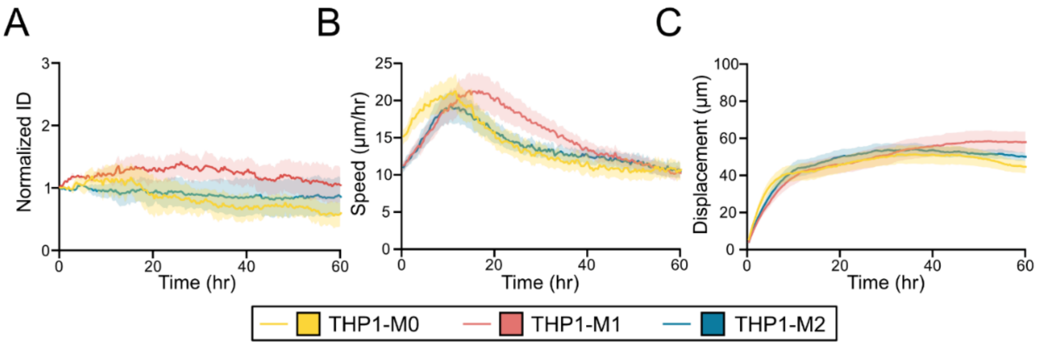


**Figure S5. Exogenous TNF-α does not cause epithelial clustering.** After adding 25ng/mL TNF-α to epithelial cell and macrophage co-cultures on soft gels, **(A)** epithelial cells remained equidistant throughout the experiment regardless of macrophage type, indicating absence of clustering. Similarly, **(B)** migration speed and **(C)** net displacement of epithelial cells remained stagnant after the first 20hr of the experiment. Line plots represent mean and shaded regions represent SEM for n = 3.
